# Contrasting patterns of interleukin signalling in LRRK2 and idiopathic Parkinson’s

**DOI:** 10.1101/2025.08.24.672005

**Authors:** Yibo Zhao, Patrick Lewis, Valentina Escott-Price, Claudia Manzoni

## Abstract

Sporadic PD and LRRK2-associated Parkinson’s Disease (PD) present with similar features although the familial form of disease is classically associated with less cognitive impairment, better REM sleep behaviour, and reduced incidence of hyposmia but more freezing gait and postural instability. Although being classified as the same disease from a clinical perspective there is an ongoing debate on whether these 2 conditions are linked to different causative molecular mechanisms. In this work we analysed the transcriptomics and proteomics profile of sporadic and LRRK2 PD patients to contrast and compare inflammatory markers in blood and CSF. Analyses showed molecular differences in the burden of inflammatory signals clearly differentiating the sporadic and the familial forms of disease. Finally, a cluster of closely connected proteins whose expression levels were changed in the familial forms of disease was highlighted. The levels of these inflammatory markers were associated with tau pathology suggesting this protein cluster might be a system model to evaluate the control of tau metabolism in the presence of LRRK2 mutations.

## INTRODUCTION

Parkinson’s disease (PD) is the second most common neurodegeneration after Alzheimer’s (AD), affecting approximately 1-3% of the population over the age of 60 (1–3). With the continued growth of the aging population, PD is anticipated to see a more than 30% increase in both prevalence and incidence by 2030. It is clinically characterised by classical motor symptoms, including bradykinesia, resting tremor, muscular rigidity, and postural instability; as well as a wide range of non-motor manifestations among which cognitive decline, autonomic dysfunctions, olfactory loss, depression, anxiety, and sleep disturbances (4). Interestingly, the non-motor symptoms can precede motor manifestations by years and significantly impact quality of life. The majority of the cases are sporadic in origin as no large effect size mutations, segregating within the families of the affected subjects, are described. These sporadic cases are supposed to originate form a cocktail of risk factors, some genetic in origin (but with small effect size), some deriving from the environment. A minority of PD cases are familial, as the disease can be traced down to a single gene mutation with large enough effect size to trigger disease. Coding mutations in the *LRRK2* gene, particularly the G2019S variant, represent the most frequent genetic cause of PD (5, 6); and are found in up to 40% of familial PD cases and up to 2% of sporadic cases, depending on the population and ethnic background studied (7–9). LRRK2 is an enzyme harbouring a kinase and a GTPase domain, multiple phosphorylation sites and protein-protein interaction domains. The physiological function of LRRK2 has not as yet been fully elucidated, however there is evidence implicating it in the control of vesicle dynamics within the endocytosis-autophagy-lysosomal axis, as well as in the orchestration of signalling downstream multiple pathways. LRRK2 function might also be tissue/cell type specific (10).

Clinically, both sporadic PD (sPD) and LRRK2-associated PD (LPD) present with similar motor features and respond well to dopaminergic therapies such as levodopa (11, 12). sPD and LPD are generally not distinguishable from each other on a case-by-case basis (13). Nevertheless, there are some important differences in the presentation of these 2 forms of disease when we compare cohorts. LPD patients classically present with less cognitive impairment or slower cognitive decline, better REM sleep behaviour, and reduced incidence of hyposmia compared to cases of sPD (13, 14). Even the motor symptoms show subtle differences with LPD cases being more prone to freezing gait and postural instability (PIGD = postural instability gait difficulty phenotype) (15–18).

The neuropathological hallmarks of PD include the degeneration of dopaminergic neurons in the substantia nigra pars compacta and the accumulation of intracellular protein aggregates of α-synuclein (Lewy bodies). Classical sPD presents with, and is defined by, α-synuclein pathology. In contrast, LPD cases can display reduced or even absent deposition of Lewy bodies; however, all LPD cases present with a certain degree of tau pathology (19, 20).

These clinical and pathological distinctions suggest that, despite the overlapping clinical features that allow for the classification of these 2 conditions under the same disease, the molecular mechanisms triggering sPD and LPD may be distinct. Our previous work has demonstrated differences in transcriptomic profiles when comparing sPD and LPD, implicating distinct biological pathways in each subtype (21). Such findings raise the important possibility that tailored therapeutic strategies — rather than a one-size-fits-all approach — may be necessary to effectively treat individuals within the heterogeneous PD population (22).

Inflammation has emerged as one of the many pathological processes implicated in PD, involving both the central and the peripheral immune systems. Although to date no specific immune-biomarker has been strongly and consistently linked to disease, elevated brain inflammation has been identified in PD postmortem brain tissues as well as peripheral immune dysfunction (23). Inflammation can be a consequence of progressive neuronal degeneration, but it is also now considered a significant contributor to PD onset and progression (24, 25) with chronic peripheral inflammation (especially localized in the gut) being linked to neurodegeneration following the gut:brain axis mechanisms (26).

LRRK2 itself is highly expressed in immune cells (27) and LRRK2 expression is upregulated in response to inflammatory stimuli (28). LRRK2 has a well-documented role in immune regulation (29) and LRRK2 variants associated with PD have been found to alter cytokine production and immune cell responses, suggesting a direct link between genetic risk of PD and immune dysfunctions (30). In the context of alterations of the immune response it is important to note that LRRK2 mutations are associate with Crohn’s disease, a type of inflammatory bowel disorder (IBD) characterized by alterations in the Th17 pro-inflammatory immune response (31, 32) and leprosy where LRRK2 has been shown to play a role in the ability of the immune system to clear pathogens engulfed by the macrophagic cell populations (33).

Using integrated multi-omics (transcriptomics and proteomics) and clinical data, in this study we addressed whether subtype-specific alterations in interleukin pathways could be detected when comparing sPD and LPD. In particular, we aimed to characterise the interleukin signalling landscape in early-stage PD, focusing specifically on comparing sporadic and LRRK2-associated forms of disease. In this respect it is worth noticing that we are not trying to define a single classical biomarker, but to develop knowledge of immune profiles (calculated as a burden of small signals in relevant pathways) that might be different across the cohorts analysed. Additionally, we evaluated whether these immune profiles correlate with clinical descriptors of disease such as indexes of cognitive function and protein pathology.

In this work we sought to improve our understanding of immune dysregulation in PD and ultimately, this work supports the broader goal of advancing subtype-specific approaches that account for the complex heterogeneity of PD.

## METHODS

### IL Pathway Protein/Gene List

The Reactome Pathway Database (*Homo sapiens*) was queried to derive a comprehensive list of protein participants for interleukins (IL) pathways (34). Specifically, we downloaded proteins annotated with all 10 child terms of “Signalling by Interleukins (R-HSA-449147)”, including “Interleukin-1 family signalling (R-HSA-446652)”, “Interleukin-2 family signalling (R-HSA-451927)”, “Interleukin-3, Interleukin-5 and GM-CSF signalling (R-HSA-512988)”, “Interleukin-4 and Interleukin-13 signalling (R-HSA-6785807)”, “Interleukin-6 family signalling (R-HSA-6783589)”, “Interleukin-7 signalling (R-HSA-1266695)”, “Interleukin-10 signalling (R-HSA-6783783)”, “Interleukin-12 family signalling (R-HSA-447115)”, “Interleukin-17 signalling (R-HSA-448424)” and “Interleukin-20 family signalling (R-HSA-8854691). Of note, the child term “Other interleukin signalling (R-HSA-449836)” was excluded due to its ambiguous classification. All entities in this paper are represented using the HUGO gene nomenclature.

### Study Cohort

Parkinson’s Progression Markers Initiative (PPMI, RRID:SCR_006431) is an ongoing observational, international, multicentred cohort study aimed at identifying the biomarkers of PD progression in a large cohort of participants (https://www.ppmi-info.org/, the current PPMI Clinical study protocol #002 is WCG approved with IRB Tracking #20200597). PPMI is a public-private partnership—is funded by The Michael J. Fox Foundation for Parkinson’s Research and funding partners, including those reported at https://www.ppmi-info.org/about-ppmi/who-we-are/studysponsors. Study protocol and manuals are available online (http://www.ppmi-info.org/study-design). PPMI enrols prodromal, recently diagnosed Parkinson’s participants and healthy controls (all after informed consent). For a more detailed cohort description please refer to https://www.ppmi-info.org/study-design/study-cohorts/. In this study, we included healthy controls, de novo PD cases without pathogenetic variations (sporadic PD or sPD), and PD cases with the LRRK2-G2019S mutation (LPD). We downloaded the participant status document (16 May 2023) and processed the sample list to remove: subjects in cohorts other than the 3 listed before; subjects whose enrolment status was recorded as “declined”, “excluded”, or “screen failed” or with additional comments such as “participants to be excluded”. Based on the enrolment status and the confirmed status (if available) healthy controls (cntrls), sPD and LPD subjects were kept. LPD subjects were further filtered to retain G2019S carriers only, leading to a total of 266 Controls, 908 sPD cases 149 LPD cases. Demographic features of subjects were derived from PPMI, including age at baseline, gender and ethnicity. Disease durations at baseline of the sPD and LPD cohort were calculated as: 𝐷𝑎𝑡𝑒 𝑜𝑓 𝑃𝑎𝑟𝑘𝑖𝑛𝑠𝑜𝑛^′^𝑠 𝑑𝑖𝑠𝑒𝑎𝑠𝑒 𝑑𝑖𝑎𝑔𝑛𝑜𝑠𝑖𝑠 − 𝐴𝑠𝑠𝑒𝑠𝑠𝑚𝑒𝑛𝑡 𝐷𝑎𝑡𝑒.

Whole-blood transcript levels and CSF protein levels of IL entities were extracted from PPMI Project 133 and Project 151 (curated) using HUGO gene symbols and Ensembl IDs on 13^th^ February 2024. Motor disability for LPD and sPD subjects were evaluated via the Hoehn and Yahr scale (H&Y). For disease outcomes, we included CSF levels of 4 biomarkers (𝛼-synuclein, A𝛽_1−42_, phosphorylated tau protein (ptau) and total tau protein (tau) and 4 cognitive scores: Montreal Cognitive Assessment (MoCA) for overall cognitive functions; the Semantic Fluency Test (SFT; total score of animal, fruit, and vegetable) for executive function; the Letter-Number Sequencing (LNS) test for executive function and working memory; and the Shared Decision Making Questionnaire (SDM). We also extracted the Cognition Status for each subject at the 3rd year visit (from baseline) as recorded by PPMI in the “Cognitive_Categorization.csv” file, which includes 5 cognition categories: “Normal”, “Cognitive Complaints”, “Mild Cognitive Impairment (MCI)”, “Dementia”, and “Indetermined”. We excluded subjects with “Indetermined” cognitive status from this study.

### Differential Expression Analyses (DEA)

Whole-blood transcript levels of IL genes were extracted and compared across the cntrls, sPD and LPD cohorts via the DESeq2 R package (35). Genes with low read counts, defined as having ≤ 15 reads in ≥ 25% of samples, were excluded (threshold adapted from the WGCNA R package) (36). Transcript level of each IL gene was compared between [cntrls vs. sPD], [cntrls vs. LPD] and [sPD vs. LPD] groups, adjusting for disease duration (scaled as required by the package for continuous variables) and sex. Naïve p-values were derived from the DESeq2 algorithm and then adjusted using the Effective Number of Tests (ENT) approach, which accounts for feature correlations. ENT was estimated via the eigenvalue-based method of Li and Ji to control the type I error rate via the R package “poolr” (37). For CSF proteins, each IL protein was evaluated using logistic regression models via the formula: 𝑃ℎ𝑒𝑛𝑜𝑡𝑦𝑝𝑒 ∼ 𝐷𝑖𝑠𝑒𝑎𝑠𝑒 𝐷𝑢𝑟𝑎𝑡𝑖𝑜𝑛 + 𝑆𝑒𝑥 + 𝐼𝐿 𝑃𝑟𝑜𝑡𝑒𝑖𝑛 𝐿𝑒𝑣𝑒𝑙. For comparisons between [cntrls vs. sPD] and [cntrls vs. LPD], cntrls were coded as 0 and cases as 1. For the comparison between the [sPD and LPD], sPD was coded as 0 and LPD as 1. p-values were adjusted using the same ENT approach as described above.

Functional enrichment via pathway over-representation was performed via the Reactome analysis tool. Results were processed as follows: i) non FDR significant terms were removed, ii) pathway coverage was calculated (reactions found/reaction total), iii) terms (pathways) containing <5 total proteins (“entities total”) (<4 in the case of sPD to retain slightly more terms for the analysis) were removed, iv) pathway terms with <10% coverage were removed, v) top term for coverage and p-value (FDR) were selected (coverage was considered when equal p-value).

### Association Analyses and 3Y Prediction of Cognitive decline

Significant transcripts from differential expression analysis (DEA) were compiled separately for the LPD and sPD cohorts. For LPD, signals were derived from [cntrls vs. LPD] and [sPD vs. LPD – adjusted for disease duration] comparisons; for sPD, signals were derived from [cntrls vs. sPD] and [sPD vs. LPD – adjusted for disease duration] comparisons selecting significant markers after multiple test correction. Of note, only signals with |log2fc| (whole-blood mRNA) or |coefficient| (CSF protein) > 60% percentile were included in further analysis. Elastic Net Regression was applied to select whole-blood transcripts or CSF proteins from the LPD and sPD lists associated with baseline cohort descriptors including: 4 CSF biomarkers (α-synuclein, Aβ₁₋₄₂, ptau, and tau) and 4 cognitive scores (MoCA, SFT, LNS, and SDM), analysed separately. Continuous variables were scaled prior input to regression models. Each model was run 100 times, and the mean coefficients across runs were used to ensure stability of feature selection. All features that were selected in at least one outcome model were then combined for downstream analysis.

Selected IL signals in the sPD and LPD cohorts were included in the logistic regression model to evaluate their contribution to predicting 3-year cognitive decline, analysed separately for each cohort. To reduce the risk of overfitting, only features with coefficients < –0.1 (negative association) or > 0.1 (positive association) were retained. Each model was adjusted for age, sex, *APOE* genotype, and disease duration at baseline. The binary outcome was defined as: 0 – normal cognitive status at 3 years; 1 – any other cognitive status at 3 years as previously described. Model performance was evaluated using 10-fold cross-validation, with the area under the receiver operating characteristic curve (AUROC) as the metric. The predictive performance of models including IL signals was compared against models with covariates only.

Elastic net regularised regression models with 10-fold cross-validation were implemented using the “glmnet” (38) and “caret” (39) packages. Receiver operating characteristic (ROC) curves were generated using the “pROC” package (40)

### Definition of the tau cluster

The IL signals that were associated with the disease predictors tau and ptau for the LPD cohort (in addition to LRRK2 and tau proteins) were used as “seeds” for downloading protein-protein interactions (PPIs) from PINOT (https://www.reading.ac.uk/bioinf/PINOT/PINOT_form.html, 02/June/2025) (41). PPIs downloaded from PINOT come from peer-reviewed literature manually annotated into PPIs databases. PPIs were quality controlled to effectively keep those with a final score (FS) equal or larger then 4 (unless one of the interaction partners was LRRK2 or *MAPT* (tau) or an interleukin marker, in which case all PPIs have been retained regardless of FS = measure of replication).

### Statistic Tools and Software

All statistical analyses were performed using R version 4.0.2 within RStudio version 2024.09.1+394. All plots were produced with ggplot2 (42). Network visualization and editing was conducted in Cytoscape version 3.8.1 (43).

## RESULTS

### Cohort QC and Characterisation

Parkinson’s disease cases, both sporadic or with the G2019S LRRK2 mutation (sPD and LPD), alongside controls (cntrls) were selected from PPMI as detailed in materials and methods. We further excluded non-white participants to minimise biases induced by population stratification (82/1321, 6.2%). Additionally, 5 cntrls exhibited H&Y score > 0 and were thereby excluded, with 235 cntrls, 856 sPD cases, 149 LPD cases remaining for the analysis. Of these, 442 subjects (132 cntrls, 246 sPD cases and 64 LPD cases) had whole-blood transcriptomics data and 659 subjects (168 cntrls, 359 sPD cases and 132 LPD cases) had CSF proteomics data available.

No significant difference was observed in age at baseline in sPD, LPD and cntrls (**Table 1**). However, the LPD cohort was composed of more females than the sPD cohort (Chi-square p = 2X10^-3^), and it showed significantly longer disease duration than the sPD cohort (Wilcoxon p-value < 1X10^-3^). Therefore, sex and disease duration were corrected (when possible) in the follow-up analyses. The sPD cohort presented significant cognition decline as measured by MoCA, SFT, LNS and SDM tests compared to cntrls (Wilcoxon p-value < 1X10^-3^). The LPD cohort also showed impaired cognitive functions but to a smaller extent, as reflected by a decreased in LNS and SDM score only (Wilcoxon p-value < 5X10^-2^ for each). Both PD cohorts showed significantly lower CSF 𝛼-synuclein, A𝛽_1−42_, tau and ptau level in comparison with cntrls at baseline (ANOVA and post hoc Tukey p-value < 1X10^-3^). Similar lower CSF Aβ_1-42_ levels were observed in the 2 PD cohorts at BL. There was no significant difference in the distribution of the *APOe4* allele across the three groups (Chi-square p = 0.896). Specifically, 25.6% of cntrls (n = 46), 24.6% of sPD cases (n = 93), and 20.7% of LPD cases (n = 30) carried at least one ε4 allele

**Table 1.** Baseline and 3 year (3Y) Characteristics of PPMI Cohort.

|  | cntrls (N = 235) | sPD (N = 856) | LPD (N = 149) |
| --- | --- | --- | --- |
| **Age (Year)** | 62.2 ± 10.9 | 63.3 ± 9.0 | 63.9 ± 7.6 |
| **Female** | 91(38.7%) | 295 (34.5%) | 75 (50.3%)^ |
| **Disease Duration (Month)** | - | 26.0 ± 24.3 | 66.5 ± 63.9^ |
| **MoCA** | 28.0 ± 1.5 | 27.0 ± 2.3* | 26.2 ± 2.9 |
| **SFT** | 52.2 ± 11.0 | 48.9 ± 11.8* | 49.9 ± 12.2 |
| **LNS** | 11.7 ± 2.8 | 11.6 ± 2.7* | 11.1 ± 3.0* |
| **SDM** | 47.0 ± 10.7 | 41.1 ± 9.8* | 41.3 ± 11.2* |
| **αSyn (pg/mL)** | 1748.3 ± 754.6 | 1518.5 ± 677.4* | 1497.1 ± 678.0* |
| **$A\beta_{1-42}$ (pg/mL)** | 1759.9 ± 828.8 | 1459.9 ± 669.2* | 1376.9 ± 372.9* |
| **total tau (pg/mL)** | 193.6 ± 79.7 | 170.4 ± 57.9* | 171.1 ± 66.1* |

| phospho tau (pg/mL) |  | 17.7 ± 8.5 | 14.9 ± 5.4* | 14.9 ± 6.0* |
| --- | --- | --- | --- | --- |
| **APOe**<br>**(N subjects)** | e2/e2 | 3 | 2 | 1 |
|  | e2/e3 | 17 | 48 | 18 |
|  | e2/e4 | 6 | 6 | 1 |
|  | e3/e3 | 113 | 235 | 96 |
|  | e3/e4 | 35 | 78 | 27 |
|  | e4/e4 | 5 | 9 | 2 |
| **3Y Cognition**<br>**Status**<br>**(N subjects)** | Normal | 142 | 205 | 64 |
|  | Cognitive Complaint | 5 | 53 | 8 |
|  | MCI | 1 | 23 | 1 |
|  | Dementia | 0 | 7 | 2 |

Abbreviations: H&Y – Hoehn and Yahr Scale; MoCA - Montreal Cognitive Assessment; SFT - Semantic Fluency Test; LNS - Letter-Number Sequencing Test; SDM - Shared Decision-Making Questionnaire; $\alpha$ Syn – $\alpha$ -Synuclein, MCI – mild cognitive impairment. \*Significant differences as compared to controls; ^Significant differences as compared to the sPD cases. Parametrical variables were compared across the 3 PPMI cohorts via ANOVA with post hoc Tukey’s test; while non-parametrical variables were compared via Wilcoxon Rank-Sum Test; categorical variables were compared via Chi-square test.

### Interleukin (IL) Pathways and Protein Composition

Reactome analysis of the “Signalling by Interleukins (R-HSA-449147)” term retrieved a total of 603 proteins implicated in 10 IL pathways. These proteins range from receptors to enzymes involved in IL processing and downstream target genes modulated by IL signalling. After removing isoforms and duplicates across IL pathways, a list of 446 unique proteins was obtained (hereafter referred to as “IL proteins”, **Table S1**). The majority of the proteins in the IL proteins list (153 and 111 proteins respectively) belonged to the IL-1 and IL-4/13 pathways. The IL-7, IL-20 and IL-6 pathways comprised the smallest number of proteins (N = 25, 25 and 24, respectively), followed by the IL-3/5, IL-10 and IL-2 pathways (N = 48, 45 and 44) and the IL-12 and IL-17 pathways (N = 56 and 72 respectively) (**Figure 1A**). A total of 341 out of 446 (76.4%) proteins were private to 1 IL pathway only, while only 6 proteins (JAK1, STAT3, JAK2, TYK2, STAT1 and JAK3) were shared across ≥ 50% of the IL pathways, suggesting that IL pathways are largely independent of each other (**Figure 1B**).

**Figure 1.**
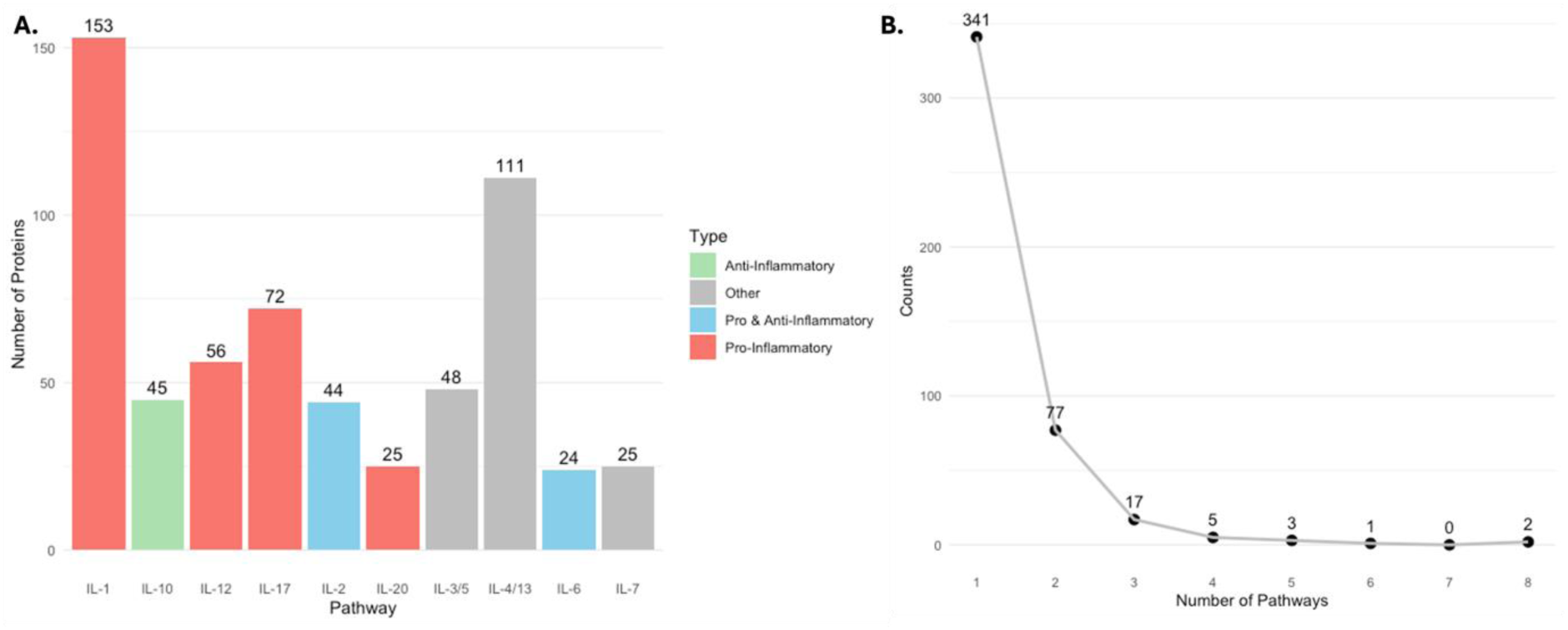
Distribution of Proteins in IL Pathways: **A.)** The bar plot shows the total number of proteins annotated for the 10 IL pathways selected from Reactome. Numbers above the bars represent the number of proteins within each pathway. **B.)** The curve represents the distribution of of protein participants across the 10 IL pathways involved in the analysis. Numbers represent the total amount of proteins shared by the number of different pathways detailed on the x-axis.

### IL Genes Presented Distinct Whole-blood Transcription Patterns in sPD vs LPD

Baseline whole-blood transcript levels of 437 out of 446 IL proteins from the IL list were derived from the PPMI database for 132 cntrls, 246 sPD cases and 64 LPD cases. Before DEA, further 89 genes were removed from analysis due to low read count. A total of 8 IL transcripts (CRK, FPR1, IL1R2, IL6R, LCP1, LMNB1, MAPK14, MCL1) presented significantly higher levels in the sPD cohort as compared to cntrls (**Figure 2A**, **Table S2**) while 46 IL transcripts were observed with significantly modified levels in the LPD cases compared to cntrls (24 transcripts downregulated and 22 upregulated, **Figure 2B**, **Table S2**). Of note, only 2 transcripts (IL1R2 and FPR1) presented the same upregulation in the 2 PD cohorts vs cntrls (**Figure 2C**). A direct comparison of the LPD cases against the sPD cohort showed 32 differentially expressed IL transcripts, including 17 upregulated and 15 downregulated signals (**Figure S1A**). After adjustment for disease duration, 20 transcripts out of 32 (62.5%; 13 upregulated and 7 downregulated) remained significant (and with the same directionality of change), suggesting that disease duration had a relatively small impact on the overall results (**Figure S1B** and **Table S2**).

**Figure 2.**
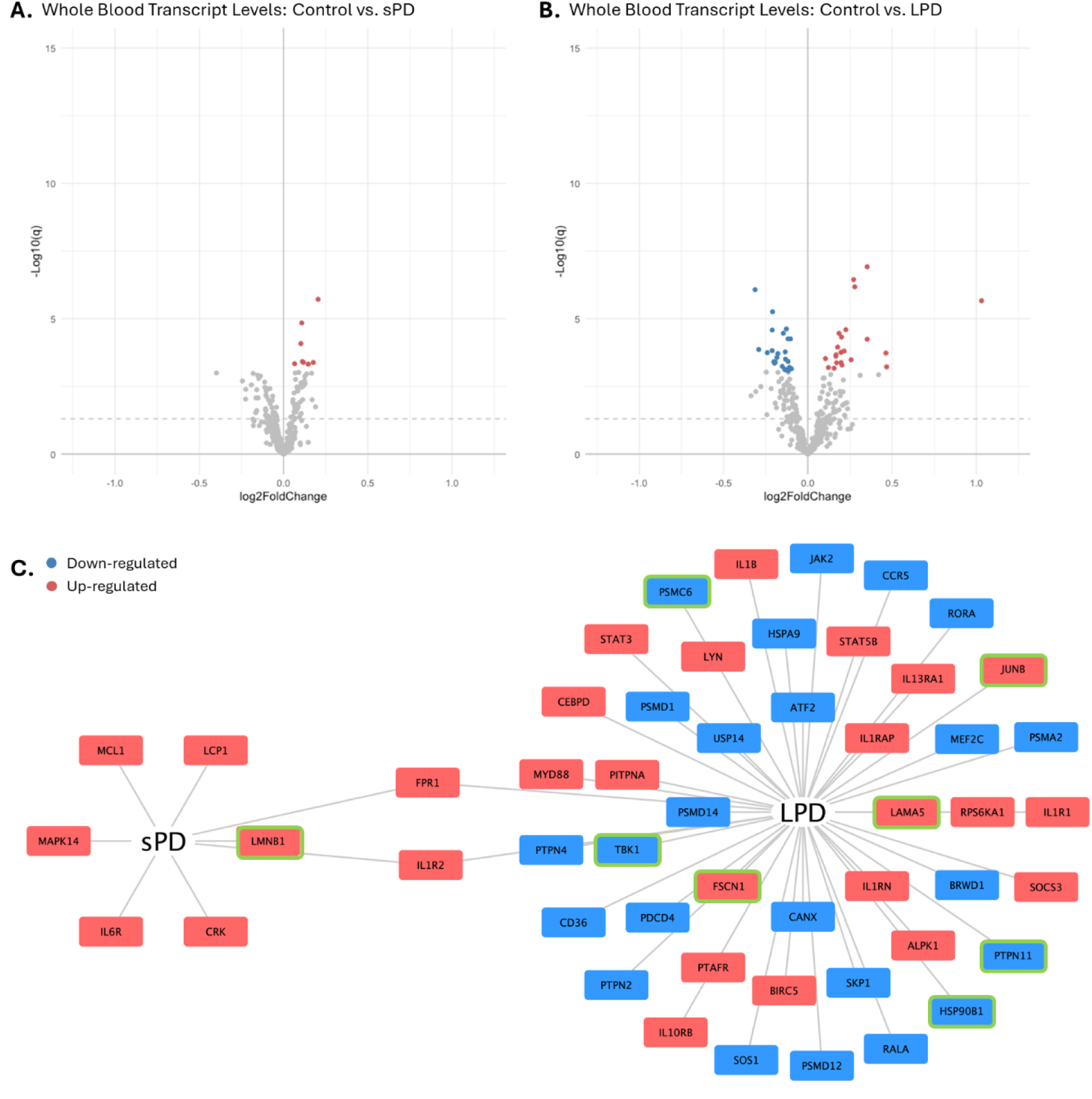
DEA on Whole-Blood Transcript Levels of IL Genes Among the sPD, LPD and Control cohorts: The volcano plots show the results of DEA on whole-blood transcript level of IL genes in the 2 PD cohorts as compared to controls; sPD in **A.)** and LPD in **B.)** Red scatters represent upregulated transcripts while blue scatters represent downregulated transcripts. The grey dash line represents unadjusted p-value threshold (p = 0.05); **C.)** The network graph shows the IL genes with significant alterations in the 2 PD cohorts as compared to controls. Red nodes represent the upregulated genes and blue nodes represent the downregulated genes. The 8 nodes with a green outline are those that remain differentially expressed when directly comparing LPD and sPD (and correcting the analysis for disease duration).

### IL CSF Proteins Presented Distinct Patterns in sPD vs LPD

Baseline CSF levels of 265 out of 446 IL proteins from the IL list were derived from PPMI database for 168 cntrls, 359 sPD cases and 132 LPD cases. Proteomics analysis showed that IL27RA and RPS27A presented significantly lower levels in the sPD cohort compared to cntrls (**Figure 3A**, **Table S3**). Of note, it is worth highlighting that IL-27 is part of the IL-12 cytokine family. The comparison between LPD cases vs cntrls showed 2 significantly decreased proteins (IL13 and PSMB6) and 5 significantly increased proteins in CSF (CXCL10, ITGB2, JAK2, LGALS9 and SOCS3; **Figure 3B**, **Table S3**). Direct comparison of the 2 PD cohorts resulted in 5 proteins (ITGB2, LGALS9, SMAD3, MAPK3 and ANXA2) with significant higher abundance and 2 proteins (IL13 and PSMB6) with significant lower levels (**Figure S2A**, **Table S3**) in LPD vs sPD. Among these signals, ITGB2, LGALS9 and IL13 remained significant after adjustment for disease duration (**Figure S2B**, **Table S3**) showing again that disease duration played a small but detectable role in differentiating the 2 cohorts of cases.

**Figure 3.**
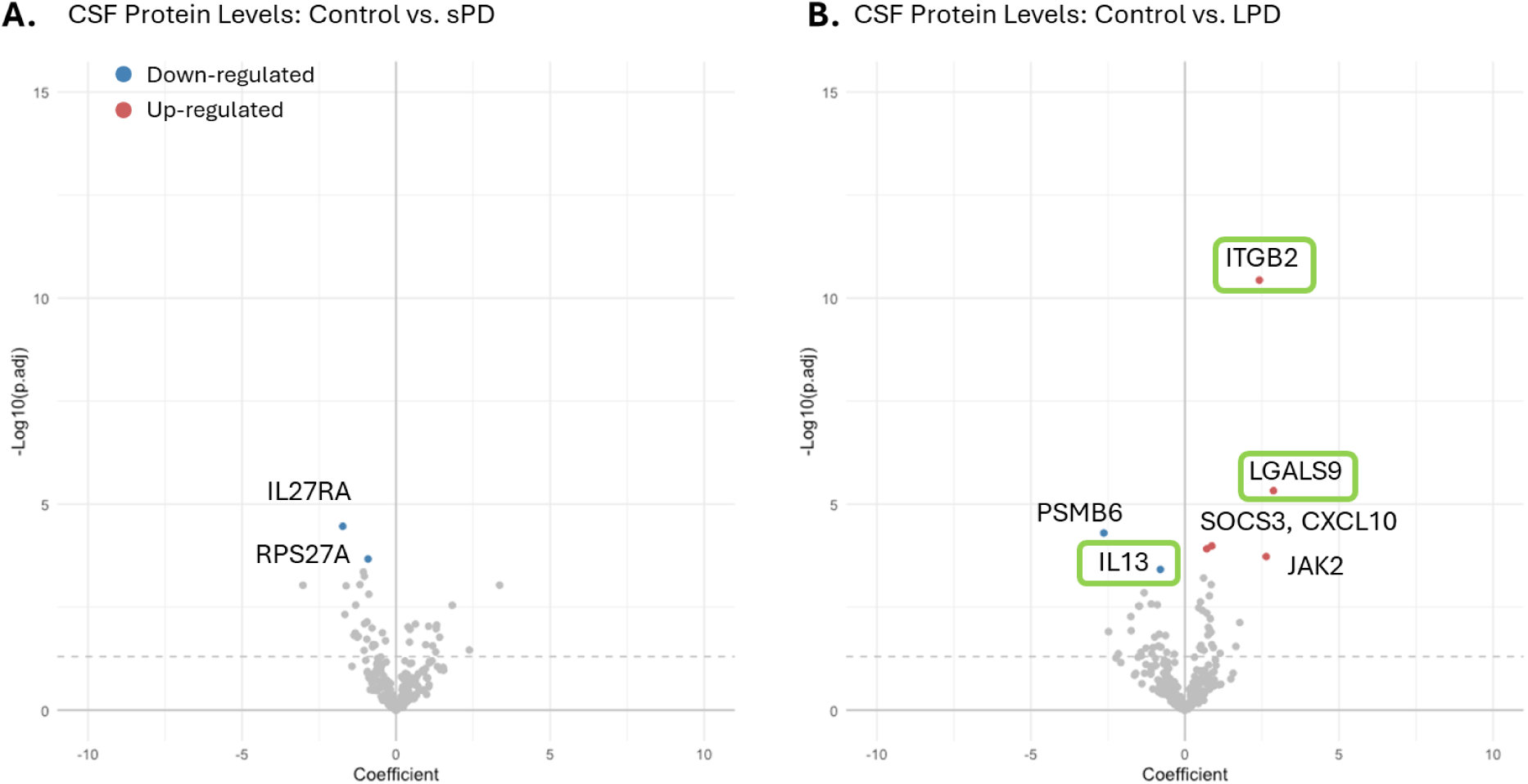
DEA on CSF Levels of IL Proteins comparing sPD, LPD and control cohorts: The volcano plots show the results of DEA on CSF levels of IL proteins in the 2 PD cohorts as compared to controls: sPD in **A.)** and LPD in **B.)** Red scatters represent upregulated proteins while blue scatters represent downregulated proteins. Nodes with a green outline are those that remain differentially expressed when directly comparing LPD and sPD and correcting the analysis for disease duration.

### Functional profiling of the differentially expressed ILs in sPD and LPD

To further evaluate the results obtained from the DEA in whole blood transcriptomics and CSF proteomics, we determined the distribution of differentially expressed signals across the 10 IL pathways. We run 1000-iteration Monte Carlo simulations by matching random lists of IL proteins (generated with the same numerosity of the DEA significant results) with the lists detailing the IL pathways composition and using cumulative distribution to determine the percentage of time the number of matches was higher or equal to the number of real matches. The analysis showed that the IL-12 pathway was borderline over-represented for the comparison [sPD vs cntrls] p = 5.1X10^-2^. On the other hand, the IL-10 pathway was over-represented (p = 7X10^-3^) while the IL-17 pathway under-represented (p = 3.9X10^-2^) for the comparison [LPD vs cntrls] (**Table 2**). In comparison, signals observed in [LPD vs sPD] (corrected for disease duration) were found with an over-representation of the IL-4/13 pathway (p = 1.3X10^-2^) and under-representation of the IL-10 pathway (p = 5X10^-2^).

**Table 2.**
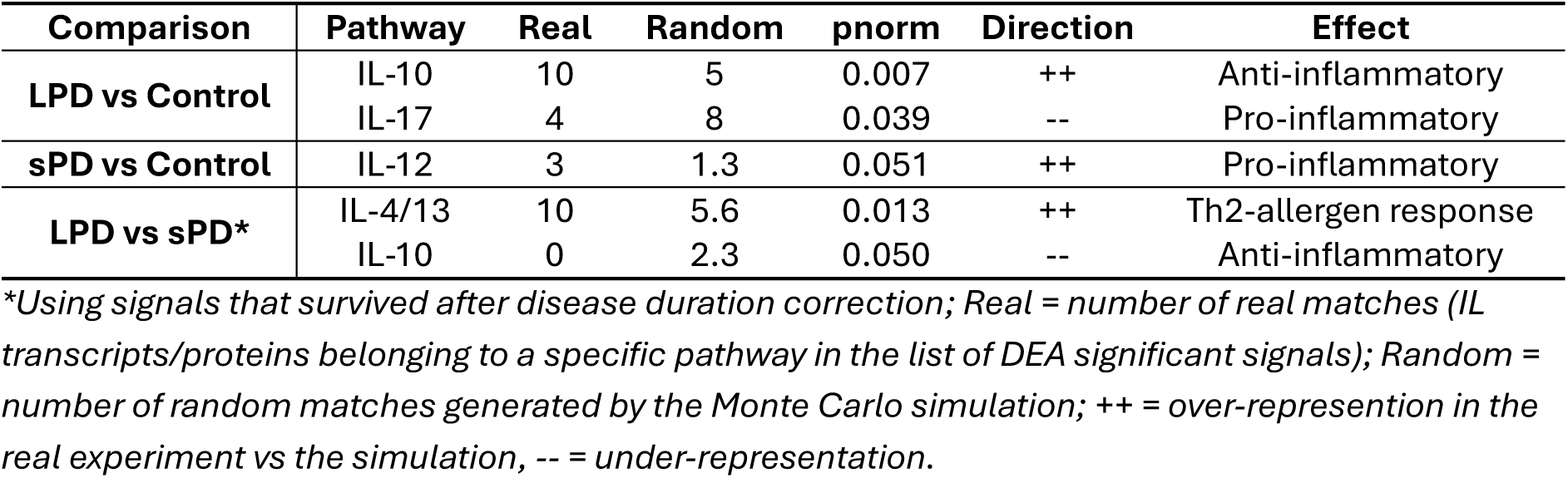
Monte Carlo Simulation Test on DEA Signals.

A second approach for functional annotation of the DEA results was conducted using the hypergeometric distribution analyses (over-representation test), performed via Reactome and corrected for multiple testing with the Benjamini-Hochberg method aimed at determine whether certain Reactome’s terms (pathway identifiers) present with a higher than random number of proteins that are common between the significant DEA results and a term themselves. Results from the over-representation tests were filtered to select the top 5 Reactome terms – pathway identifiers – based on significance (p-value) and on pathway coverage which takes into consideration in how many “units” or steps of the pathway, the proteins in the significant lists out of DEA feature **(Table 3**). Results showed a pleiotropy of pathways (IL-4, IL-13, IL-10, IL-17, IL-6 and IL-1) to be associated with the LPD scenario, while the IL-12 pathway was again prioritised solely and specifically in the enrichment of DEA signals from the comparison [sPD vs cntrls] both in terms of adjusted enrichment p-value (FDR) and pathway coverage.

**Table 3.**
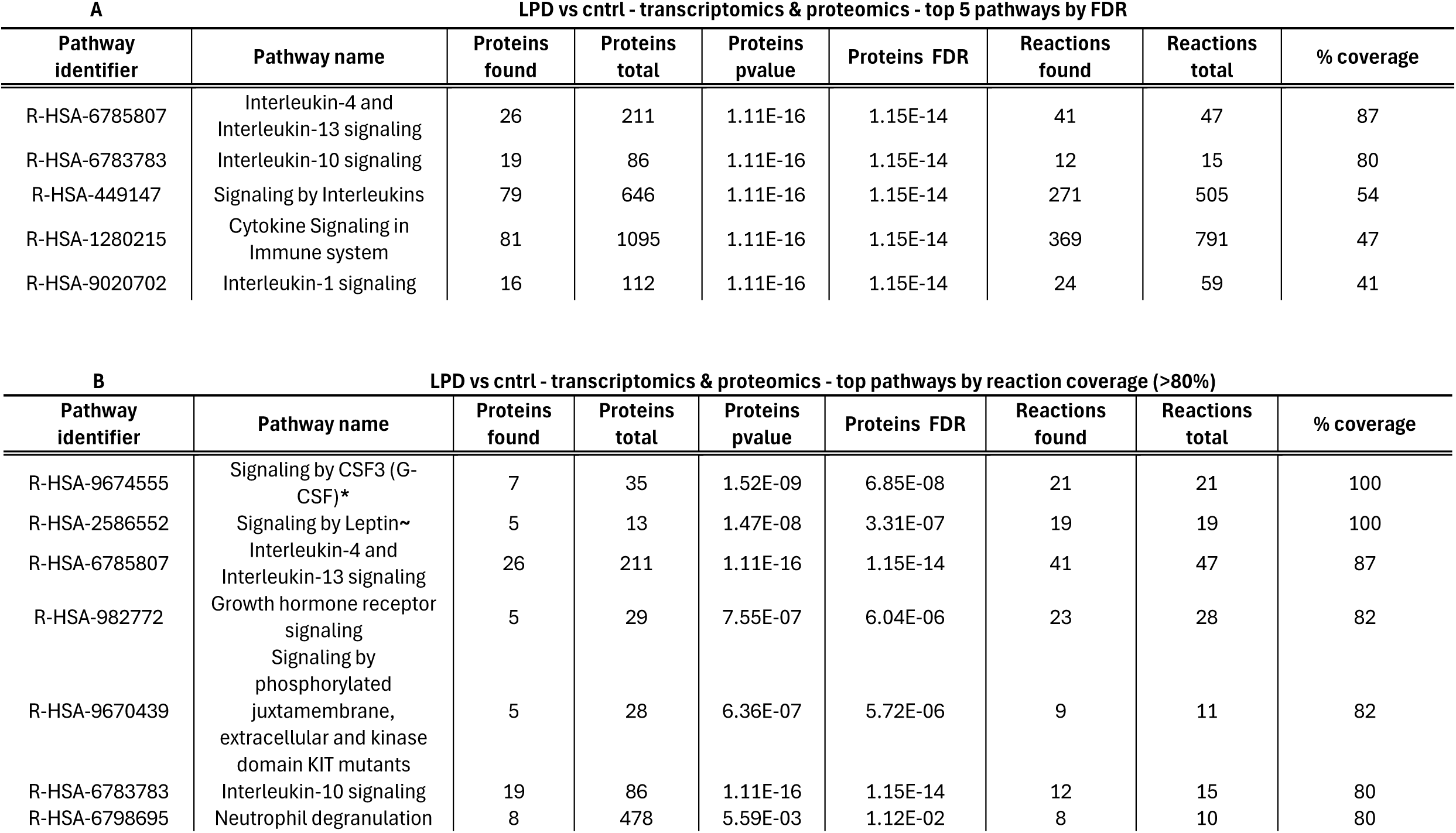
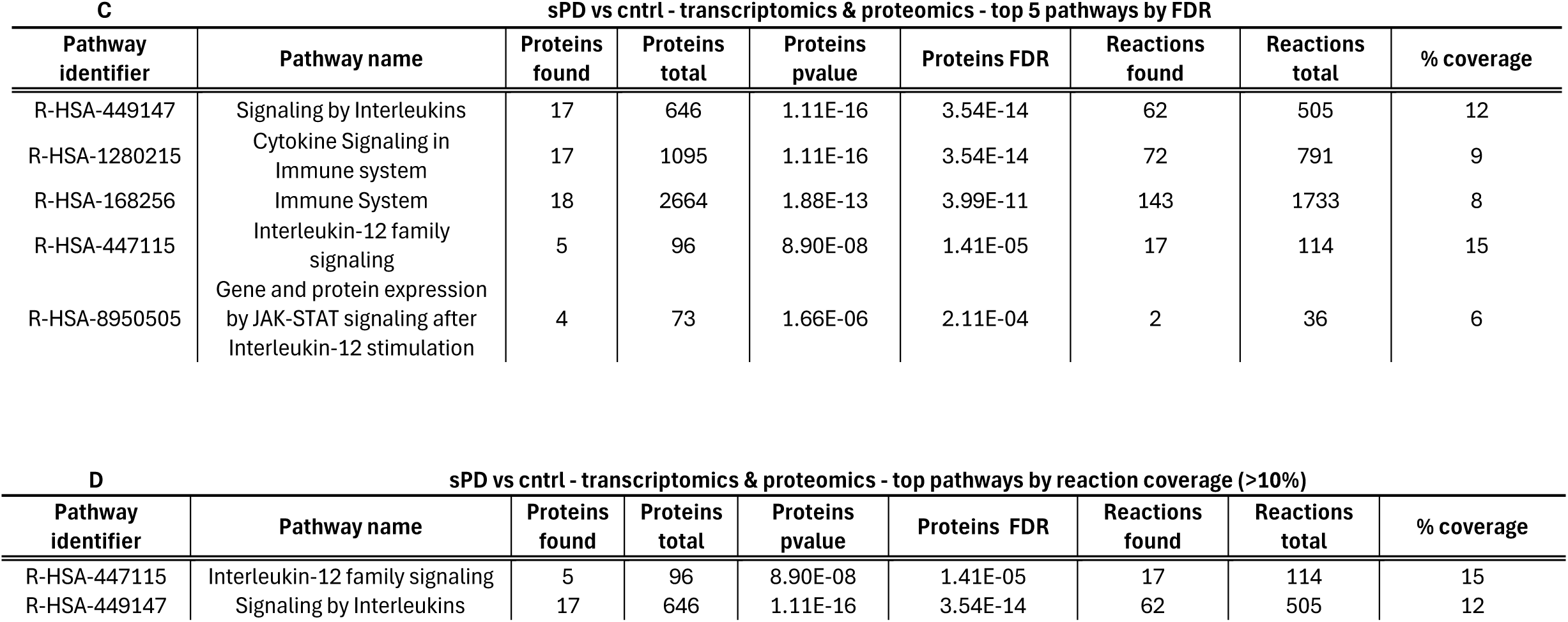
**Hypergeometric Distribution Analyses on DEA Signals** - Table **3A** and **3C** report the top 5 pathway terms by p-value out of the over-representation analysis (hypergeometric distribution) test performed with the significant DEA signals. **Table 3B** and **3C** show the top pathway terms by coverage calculated based on the number of reactions (pathway steps). * granulocyte colony-stimulating factor (CSF3) signalling promotes production & differentiation of neutrophils and granulocytes; ∼Leptin is a hormone secreted by adipocytes able to modulate the inflammatory response.

### IL alterations in transcription and expression are associated with disease descriptors of protein pathology and cognitive decline

In the case of sPD, the combination of transcripts and proteins significantly altered in [sPD vs. cntrls] and [sPD vs. LPD] (after correction for disease duration) provided a total of 48 signals to be used for the adjusted Elastic Net regression model to determine association with disease descriptors related to protein pathology and cognitive decline (**Figure 4A**, **Table 4A**, **Table S5**). We found that CSF 𝛼-synuclein was correlated with CSF levels of LGALS9 and ITGB2 (mean coefficients = 0.6 and 0.11, respectively) as well as with whole blood RNA levels of JUNB and IL17RC (mean coefficients = 0.1 and −0.07, respectively). CSF A𝛽_1−42_ was associated with whole-blood transcript levels of LCN2 and IL6R (mean coefficients = 0.08 and −0.06) as well as CSF levels of RPS27A, LGALS9 and ITGB2 (mean coefficients = −0.06, 0.34 and 0.14). Both CSF ptau and total tau levels related to CSF level of LGALS9 (mean coefficients = 0.64 and 0.65) while CSF levels of ITGB2 correlated with levels of ptau only (mean coefficient = 0.07). In terms of cognition scores, SFT showed correlation with whole-blood transcript levels of MCL1 and CSF levels of LGALS9 (mean coefficients = 0.08 and −0.07 respectively); MoCA scores correlated with CSF levels of LGALS9 only (mean coefficients = −0.1).

**Figure 4.**
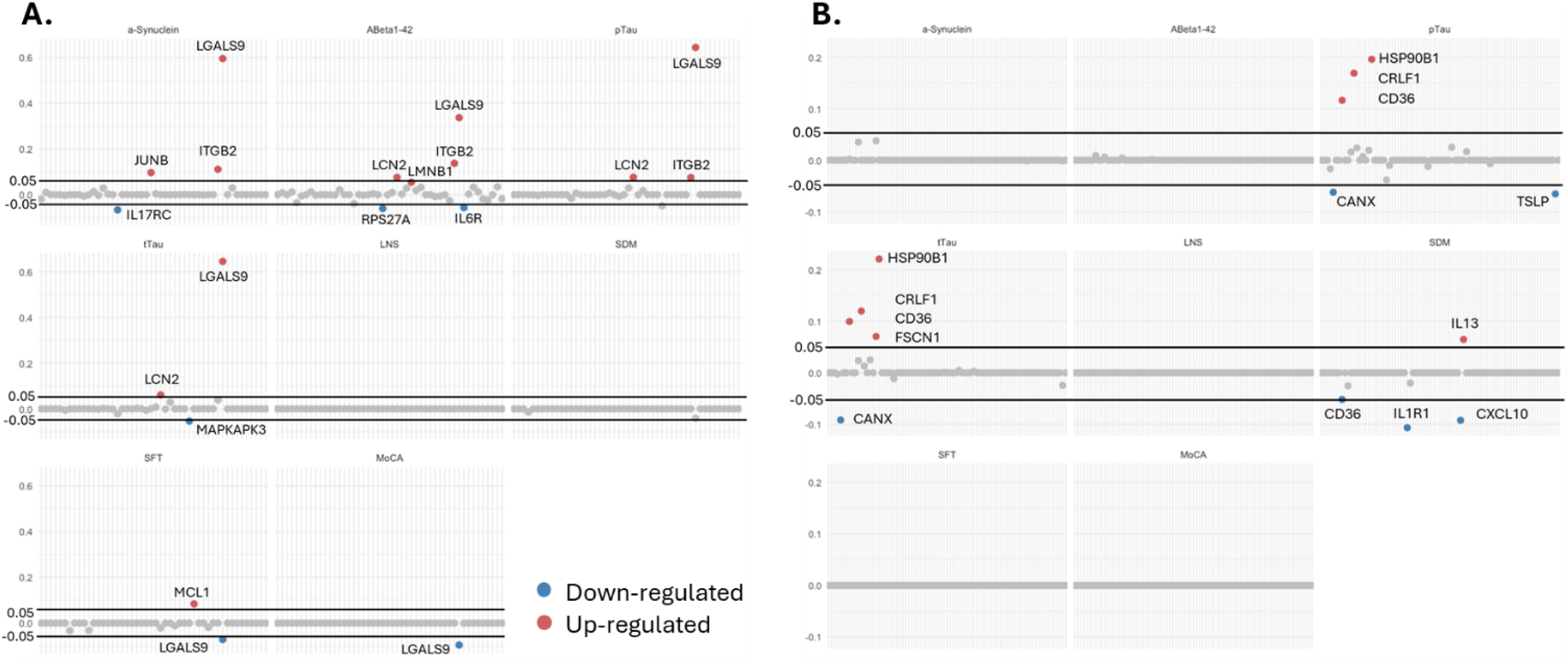
Markers in IL pathway association with disease descriptors: Dot plots show IL-pathway-related features selected by Elastic Net Regression across nine cognitive outcomes at 3Y, including CSF biomarkers (α-synuclein, Aβ₁₋₄₂, ptau, and total tau) and cognitive scores (LNS, SDM, SFT, and MoCA). Each elastic net model was repeated 100 times, and the mean coefficient across runs was used to assess feature stability. Red dots indicate features with a mean coefficient > 0.1, and blue dots indicate features with a mean coefficient < −0.1. Each panel corresponds to a specific cognitive outcome. **A)** sPD, **B)** LPD.

**Table 4.** Significant associations after Elastic Net Regression. Mean coefficients after Elastic Net Regression are reported, both protein levels in the CSF (p) and transcript levels in whole blood (t) are considered.

| marker | sPD |  |  |  |  |  | marker | LPD |  |  |
| --- | --- | --- | --- | --- | --- | --- | --- | --- | --- | --- |
| | α-syn | Aβ<sub>1-42</sub> | ptau | tau | SFT | MoCA | | ptau | tau | SDM |
| LGALS9 (p) | 0.6 | 0.34 | 0.64 | 0.65 | -0.07 | -0.10 | HSP90B1 (t) | 0.21 | 0.23 |  |
| ITGB2 (p) | 0.11 | 0.14 | 0.07 |  |  |  | CRLF1 (t) | 0.18 | 0.12 |  |
| JUNB (t) | 0.1 |  |  |  |  |  | CD36 (t) | 0.12 | 0.10 | -0.06 |
| IL17RC (t) | -0.07 |  |  |  |  |  | CANX (t) | -0.07 | -0.09 |  |
| LCN2 (t) |  | 0.08 | 0.08 | 0.06 |  |  | TSLP (t) | -0.07 |  |  |
| RPS27A (p) |  | -0.06 |  |  |  |  | FSCN1 (t) | 0.07 |  |  |
| IL6R (t) |  | -0.06 |  |  |  |  | IL13 (p) |  |  | 0.07 |
| LMNB1 (t) |  | 0.05 |  |  |  |  | CXCL10 (p) |  |  | -0.10 |
| MAPKAPK3 (t) |  |  |  | -0.05 |  |  | IL1R1 (t) |  |  | -0.11 |
| MCL1 (t) |  |  |  |  | 0.08 |  |  |  |  |  |

For LPD, the combination of transcripts and protein levels alterations in [LPD vs cntrls] and [LPD vs sPD] (after correction for disease duration) provided a total of 81 signals to be used for the adjusted Elastic Net regression model (**Figure 4B**, **Table 4B, Table S5**). Among these, the majority of the associations were with CSF tau and ptau comprising: whole-blood transcript levels of HSP90B1 (mean coefficients = 0.21 and 0.23 for ptau and total tau respectively), CRLF1 (mean coefficients = 0.18 and 0.12 for ptau and total tau respectively), CD36 (mean coefficients = 0.12 and 0.10 for ptau and total tau respectively), CANX (mean coefficients = −0.07 and −0.09 for ptau and total tau respectively). Whole-blood transcript levels of FSCN1 and CSF protein levels of TSLP were associated with ptau levels only (mean coefficients = 0.07, −0.07 respectively). CSF CXCL10 and IL13 protein and whole-blood IL1R1 and CDC36 transcript levels were related modulation of baseline SDM score at recruitment (mean coefficients = −0.1, 0.07, −0.11 and −0.06).

### IL markers associated with cognitive decline and protein pathology in LPD can predict 3Y cognitive decline

The IL markers associated with cognitive decline and protein pathology in LPD (HSP90B1, CRLF1, CD36, CANX, TSLP, FSCN1, IL13, CXCL10 and IL1R1; N = 9) were included in a logistic regression model to assess their performance in improving the prediction of cognitive decline (M_IL_) at 3 year time as compared to a simple prediction model (M_ref_) built only with demographic descriptors (including age, gender, *APOe* genotype and disease duration - M_ref_). Ten-fold cross-validation showed that IL predictors improved the performance of the prediction model, shifting the AUROC: M_ref_ = 0.6836 to AUROC: M_IL_ = 0.8918 (**Figure 5A**). In the sPD scenario where the use of IL predictors significantly associated with cognitive decline and protein pathology (LGALS9, ITGB2, JUNB, IL17RC, LCN2, RPS27A, IL6R, LMNB1, MAPKAPK3 and MCL1; N = 10) IL predictors did not consideably improve the prediction of cognitive decline over 3 years: AUROC: M_ref_ = 0.6700 vs AUROC:M_IL_ = 0.7036 (**Figure 5B**).

**Figure 5.**
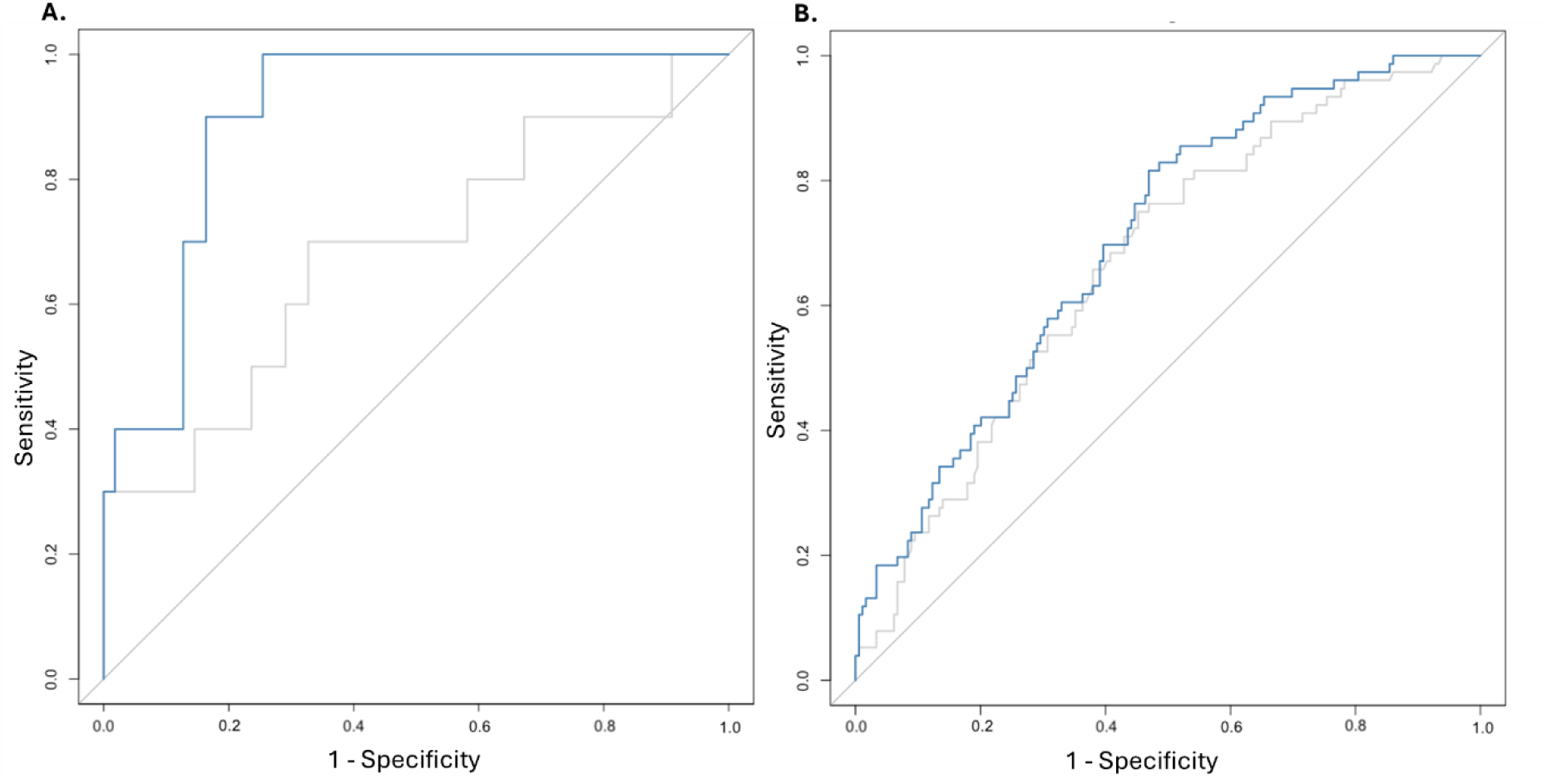
Alterations in IL pathway predicts 3 years cognitive decline in the LPD cohort. ROC curves showing the predictive performance of IL-pathway-related features for 3-year cognitive decline in the LPD cohort (**A**) and sPD (**B**). Predictive models were adjusted for age, gender, APOe genotype, and disease duration, and evaluated using 10-fold cross-validation.

### Identification of the cluster of proteins involved in tau modulation in LPD

Considering that tau pathology is highly relevant for LPD (but not for sPD), it was interesting to note that 6 out of 9 IL markers (67%) in LPD (while only 2 out 10 markers in sPD – 20%) were associated with levels of tau/ptau. We postulated that this might provide a starting point for developing a systems biology model for the molecular process behind the altered metabolism of tau in LPD. We therefore considered the differentially expressed transcripts in LPD that correlated with CSF tau/ptau levels and extracted the protein-protein interaction network around them including direct and indirect (via 1 bridge) protein interactions between: the IL markers, LRRK2 and/or tau. Direct protein interactions were considered regardless of the confidence score while indirect interactions were capped to high confidence score only.

We found that 4 out of the 6 markers analysed were connected in a protein cluster comprising both LRRK2 and the protein tau (*MAPT* gene) and containing both direct interaction across these proteins as well as indirect interaction mediated via 11 bridges. The 4 IL markers were connected directly (CANX:HSP90B1); via communal bridge proteins (CD36:APP:CANX); or via LRRK2 and tau (CANX:LRRK2:FSCN1:MAPT:CANX and HSP90B1:MAPT:FSCN1:LRRK2:HSP90B1). A topological analysis of the connection between LRRK2 and tau highlighted the relevance of the IL markers CANX, HSP90B1 and FSCN1 which effectively functioned as bridges crossing LRRK2 and tau (**Figure 6**). Another relevant protein for the network connectivity was 14-3-3γ (*YWHAG* gene) that was added to the cluster as direct protein interactor of LRRK2, tau and the IL marker HSP90B1.

**Figure 6.**
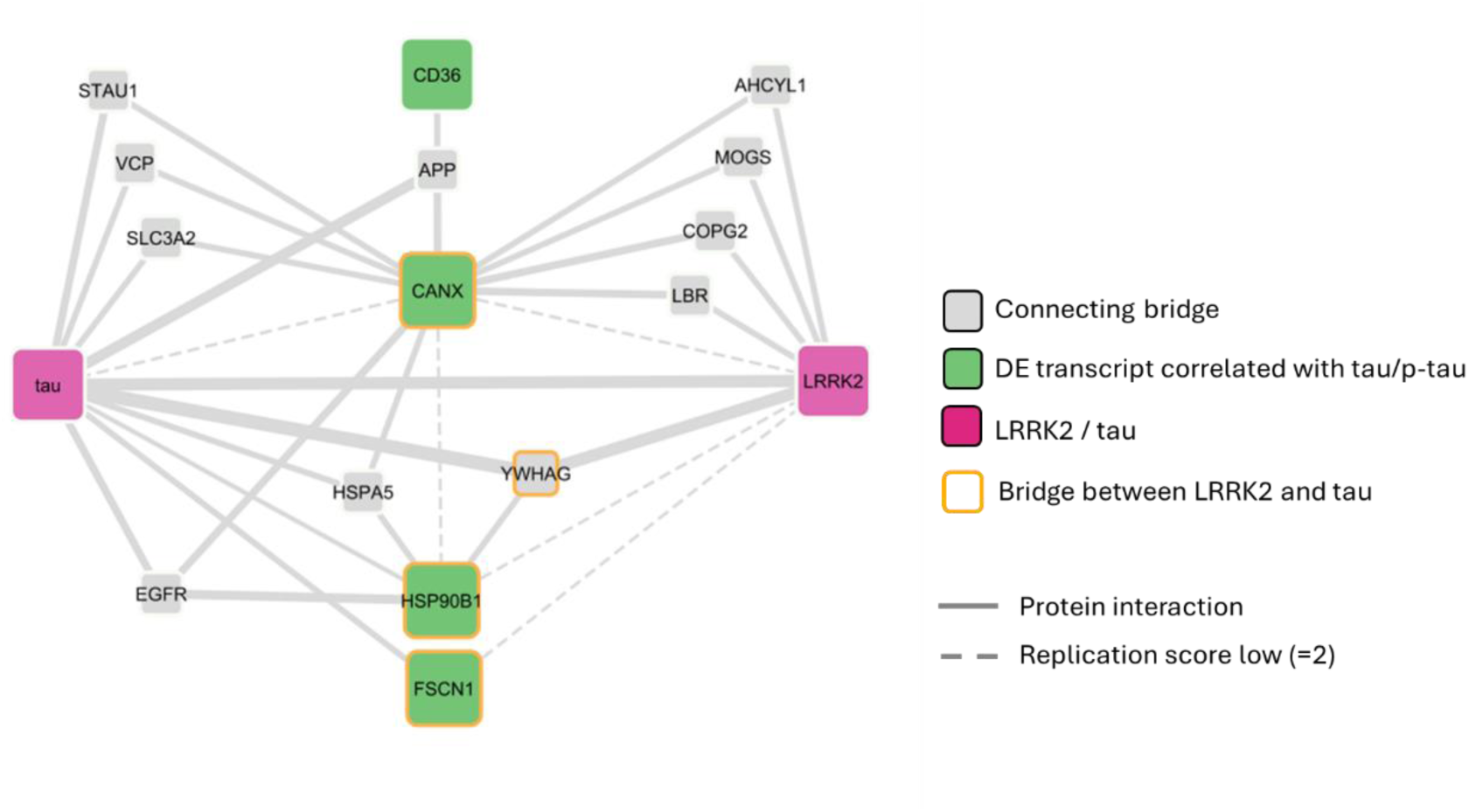
The tau cluster: The network graph shows the molecular interactions among IL predictors for tau/p-tau levels in the context of LPD. Interaction types include Protein-Protein interactions represented as solid lines whose thickness is directly proportional to the replication score; dotted line represent a non-reproduced interaction, reported only 1 time in literature (1 publication and 1 interaction detection method only).

## DISCUSSION

LRRK2 genetic mutations are strong risk factors for the triggering of PD, presenting with a clinical phenotype very similar to that of the late-onset sporadic disease. Some differences do exist in the clinical presentation between the 2 forms of PD i.e. the PIGD phenotype is generally less prominent in sPD vs LPD, while cognitive decline is less impactful and alteration of REM sleep less prevalent in the genetic disease (20). But in general, and differently from other genetic forms of Parkinsonism, LPD is considered a milder, but closely resembling version of sPD from a clinical perspective. From a pathological point of view, one very important discriminator between sPD and LPD is the deposition of Lewy bodies. While sPD is a synucleinopathy, with almost all cases being positive for α-synuclein pathology in the form of Lewy bodies, only ∼60% of LPD cases present with α -synuclein alterations, while all of them show a degree of tau proteinopathy (44). These considerations have given rise to the still unsolved conundrum of whether sPD and LPD – albeit being characterized by a similar clinical phenotype – might derive from distinct molecular alterations; a question of primary importance when it comes to drug development and clinical trials for PD.

One of the open questions within the PD field is whether the molecular mechanisms at play behind sporadic and genetic forms of PD might indeed be partially distinct. Here we analysed the cytokine environment (both in terms of transcriptomics and proteomics) in samples derived from whole blood or CSF of PD patients to define an immune signature at differential expression level. First, we isolated significant signals for transcripts (or proteins) in the interleukin pathways that are differentially expressed between controls and cases as well as sPD vs LPD. It is worth noticing that although significant, these signals have a small FC which we accepted as we are not trying to define a single classical biomarker, but to develop knowledge of immune profiles; in other words, we are more interested in the bulk effect rather than in the individual signals. Additionally, we are here testing differential expression levels, we are not directly accessing interleukin levels in serum/plasma as done by the majority of the biomarker studies available in literature. The results showed that the burden of immune signals calculated for sPD and LPD were intrinsically different with LPD featuring alterations in the anti-inflammatory machinery while sPD being associated with a burden of inflammatory signals, specifically belonging to the (extended) IL-12 pathway.

The LPD scenario was enriched for the anti-inflammatory signals typical of the IL-10 pathway, the pro-inflammatory IL-17 pathway showed under-representation while the IL-4/13 pathways were overrepresented. IL-10 is a key immunosuppressant molecule, regulator of the immune response involved in hampering inflammation during infection, thus reducing potential immune-response induced damage (45). It also directly suppresses inflammatory Th17 cells (46). Both IL-4 and IL-13 are crucial for the Th2 immune response against extracellular pathogens and allergens, in the control of excessive inflammation via modulation of Th1 responses that could harm tissue homeostasis, as well as in the induction of functional changes in epithelial and endothelial cells during inflammation and tissue repair (47, 48). In order to preserve homeostasis, inflammatory responses are balanced with anti-inflammatory feedbacks, typical of the Th2 immune response (interleukins 4, 5, 13 and 10 based), preventing excessive damage and the triggering of autoimmune attacks (49). LPD showed association with granulocyte colony-stimulating factor (CSF3) and leptin signalling the first involved in the control of neutrophils and granulocytes and the second being able to modulate the inflammatory response. Considered holistically, our data suggest the cytokine scenario for LPD (at the periphery and CSF) to be that of increased anti-inflammatory feedbacks with a potential modulation of the innate immune system that might be triggered by chronic inflammation due to the constant neurodegeneration in the presence of the G2019S LRRK2 mutation.

The role of LRRK2 within the immune system has not been clearly defined, and there is evidence for a dichotomy existing between central and peripherical functions (50). While central activation of LRRK2 in microglia and astrocytes is generally believed to results in an increased inflammatory response with consistent cytokine production (51); the effect of LRRK2 activity in the peripheral immune system is less clear. However, and in line with our findings, a body of literature exists suggesting LRRK2 hyperactivation to be linked with a reduction of inflammation at the periphery while the contrary, i.e. removal of LRRK2 kinase activity (or the entire LRRK2 protein) to be associated with reduce anti-inflammatory feedbacks. For example, in macrophages, the LRRK2 G2019S mutation did not increase the secretion of LPS-driven inflammatory cytokines (52); however, the removal of LRRK2 via knock-out (KO) seemed to reduce the anti-inflammatory status as stimulation of RAW264.7 murine macrophages (LRRK2 KO lines) showed reduced IL-10 release (53). On a similar note, LRRK2 KO or LRRK2 kinase inhibition in mouse bone marrow derived macrophages reduced IL-10 secretion after infection with M. tuberculosis, while G2019S knock-in (KI) increased secretion of IL-10 (54). A missense variant in LRRK2 (M2397T) that modulates protein turnover by substantially reducing LRRK2 half-life - and therefore reducing LRRK2 levels (55) – has been associated with increased inflammation via Th1 response in leprosy patients (56); while the LRRK2 R1628P variant (possibly associated with LRRK2 gain of kinase function) was found to be protective by reducing Th1 response in the same disease (57).

The sPD cases were associated with fewer significant expression changes, thus the burden analysis that was conducted in this study was less powerful. However, and in contrast to LPD, sPD showed a more simple, pro-inflammatory profile with over-representation of the IL-12 pathway, typically linked to the Th1 immune response. The IL-12 pathway includes IL-23, IL35 and IL-27 and all these related cytokines have a pro-inflammatory role (58).

The extant literature describing the immune system in both sPD and LPD is complex, with conflicting reports of inflammatory profiles. However, our results are consistent with the literature showing a contrasting behaviour of the peripheral immune system comparing sPD and LPD. sPD has been found associated with pro-inflammatory signals while the peripheral inflammation of LPD was shown to be similar to controls. For example, Munoz-Delgado et al. (24) evaluated peripheral immune cells showing that patients with sPD had alterations in lymphocytes and neutrophils resulting in a higher neutrophil-to-lymphocyte ratio; this was interpreted as marker of systemic inflammation. The same inflammatory response was not found in LPD where both lymphocytes and neutrophils were similar to controls. Thaler at al. (59) did not find any difference in the protein levels of several cytokines (in peripheral blood and CSF) when investigating LPD concluding no evidence of dysregulation of pro-inflammatory signals was found in LPD. On the other hand, sPD has been linked to an increased inflammatory profile both at the periphery and centrally. Many cytokines have been evaluated but Rentzos et al. (60) who found the ratio of IL-12-p40 and IL-10 to be altered in the serum of sPD patients leading to a more pro-inflammatory environment. This alteration in the IL-12/10 ratio was also replicated in Brockmann at al. (61) when analysing sPD, while the same was not identified in LPD that presented with a control-like immune profile.

In summary, and in accordance with some of the literature, with our model (based on the analysis of blood transcriptomics and CSF proteomics) we showed that the peripheral immune profile of LPD might be associated with increased anti-inflammatory feedbacks mainly involving IL-10 and IL-4/13 pathways. This was in contrast with sPD that presented with increased inflammation associated with IL-12 pathways.

It has been shown that inflammatory markers and in general the inflammatory status might correlate with disease progression or subtyping. For example increased BDNF in serum has been shown to correlate with a more severe motor/non motor presentation in LPD (61); high concentrations of IL-6 was associated with greater Aβ deposition and mild cognitive impairment in a cohort of neurological healthy older adults (62); an inflammatory finger print has been calculate to predict clinical outcome in patients with frontotemporal dementia (FTD) (63). We therefore verified whether in our cohorts the differentially expressed inflammatory markers could be associated to disease descriptors defining both the cognitive status (i.e. MoCa, SFT, LNS, and SDM tests) and the alteration in protein homeostasis (Aβ, tau, p-tau). We isolated a number of markers associated with the disease descriptors and verified that they contribute to the prediction of cognitive decline of the LPD cohort over a 3 years period, as compared to a simpler prediction model which only used demographic descriptors including age, gender, *APOe* genotype and disease duration.

In LPD, 6 out of the 9 IL markers associated with the disease descriptors (HSP90B1, CRLF1, CD36, CANX, TSLP, FSCN1) were significantly linked with levels of tau/p-tau specifically. We considered this to be an interesting result since LPD but not sPD is linked to tau proteinopaty. Therefore, we draw data from peer-review literature to design the protein interaction network around these markers focusing on the overlaps with the LRRK2 and tau (*MAPT*) interactomes. Interestingly we found that protein interactions have been reported across these proteins thus defining of a connected unit or local cluster with the principal components being LRRK2, tau, FSCN1, CANX, HSP90B1 and 14-3-3 gamma. Proteins do interact when they share functions or are involved in the same process, therefore it is possible that this local cluster holds functional information related to the crosstalk of LRRK2 and tau in the context of inflammation in LPD.

The tau protein and LRRK2 are connected directly but also via FSCN1, CANX, HSP90B1 and 14-3-3 gamma. Fascin actin-bundling protein 1 (FSCN1) is an actin-binding protein that organizes F-actin into parallel bundles within the cytoskeleton responsible for the formation of cell projections comprising the axon growth cone, while playing an important role in cell adhesion and migration (64). Interestingly, fascin has been identified as tau interactor in a study focused on endoplasmic reticulum (ER) proteins (65). We found FSCN1 altered in expression levels when comparing the blood transcriptomics profiles of [cntrl vs LPD] and [sPD vs LPD]. Calnexin (CANX) is a calcium-binding protein that plays a role in ER quality control - specifically for glycoproteins (66), as well as receptor trafficking and mediated endocytosis in neuronal cells. We found CANX altered in expression levels when comparing the blood transcriptomics profiles of [cntrl vs LPD]. Endoplasmin (HSP90B1) is a chaperon protein involved in the folding and trafficking of proteins in the ER. We found HSP90B1 altered in expression levels when comparing the blood transcriptomics profiles of [cntrl vs LPD] and [sPD vs LPD]. 14-3-3γ (*YWHAG* gene) is another strategic component of the LRRK2/tau cluster from a topological perspective as it bridges LRRK2 and tau, while connecting to HSP90B1. 14-3-3 proteins have been shown to take part into the shuffling of proteins between the ER and the plasma membrane (67). Unfortunately, many interactions within this local cluster have been reported only via high throughput assays and therefore they lack low-scale validation (65, 68–70). However, if these interactions hold true they might suggest an interplay of tau and LRRK2 at the ER via a combination of activities modulated by fascin, calnexin and endoplasmin. We observed an expression change of these proteins that correlated with measures of tau pathology in LPD patients. We therefore hypothesised that this protein unit might be able, under physiological conditions, to control the metabolism and phosphorylation of tau. Upon mutation of LRRK2, the entire unit might become dysfunctional, with measurable changes in expression levels, and this in turn might contribute to the shift in the phosphorylation and/or total amount of tau to a pathological level as observed in LPD but not in sPD.

In conclusion, this work highlights subtle but consisted molecular differences related to interleukin signalling between the sporadic and the familial (LRRK2) forms of PD, suggesting the existence of discrepancies in the underling mechanisms of disease that potentially have an impact on research focused on biomarker discovery and therapeutics development. Additionally, this suggests that the inflammatory profile should probably be taken into consideration when considering patient stratification. Finally, we highlighted a cluster of closely connected proteins that mediates the interplay between LRRK2 and tau and whose expression levels are altered in LPD in association with tau pathology. Albeit further functional validation is mandatory, we suggest this protein cluster might contribute to the control of tau metabolism in the presence of LRRK2 mutations.

